# Human bone marrow disorders recapitulated in vitro using organ chip technology

**DOI:** 10.1101/458935

**Authors:** David B. Chou, Viktoras Frismantas, Yuka Milton, Rhiannon David, Petar Pop-Damkov, Douglas Ferguson, Alexander MacDonald, Özge Vargel Bölükbaşı, Cailin E. Joyce, Liliana S. Moreira Teixeira, Arianna Rech, Amanda Jiang, Elizabeth Calamari, Sasan Jalili-Firoozinezhad, Carlos F. Ng, Youngjae Choe, Susan Clauson, Kasiani Myers, Robert P. Hasserjian, Richard Novak, Oren Levy, Rachelle Prantil-Baun, Carl D. Novina, Akiko Shimamura, Lorna Ewart, Donald E. Ingber

**Affiliations:** Wyss Institute for Biologically Inspired Engineering at Harvard University, Boston, MA 02115, USA; Department of Pathology, Massachusetts General Hospital, Boston, MA 02115, USA; AstraZeneca, Drug Safety and Metabolism, IMED Biotech Unit, Cambridge, UK; Dana Farber/Boston Children’s Cancer and Blood Disorders Center, Boston, MA 02115, USA; Department of Cancer Immunology and Virology, Dana-Farber Cancer Institute, Boston, MA 02115, USA; Department of Medicine, Harvard Medical School, Boston, MA 02115, USA; Department of Bioengineering and iBB ‐ Institute for Bioengineering and Biosciences, Instituto Superior Técnico, Universidade de Lisboa, Lisboa, Portugal; Division of Bone Marrow Transplantation and Immune Deficiency, Cincinnati Children’s Hospital, Cincinnati, OH 45229, USA; Broad Institute of Harvard and MIT, Cambridge, MA 02141, USA; Vascular Biology Program and Department of Surgery, Children’s Hospital and Harvard Medical School, Boston, MA 02115, USA; Harvard John A. Paulson School of Engineering and Applied Sciences, Cambridge, MA 02139, USA

**Keywords:** hematopoiesis, microfluidic, organ-on-chip, myelosuppression, radiation, Shwachman-Diamond Syndrome

## Abstract

Understanding human bone marrow (BM) pathophysiology in the context of myelotoxic stress induced by drugs, radiation, or genetic mutations is of critical importance in clinical medicine. However, study of these dynamic cellular responses is hampered by the inaccessibility of living BM *in vivo*. Here, we describe a vascularized human Bone Marrow-on-a-Chip (BM Chip) microfluidic culture device for modeling bone marrow function and disease states. The BM Chip is comprised of a fluidic channel filled with a fibrin gel in which patient-derived CD34+ cells and bone marrow-derived stromal cells (BMSCs) are co-cultured, which is separated by a porous membrane from a parallel fluidic channel lined by human vascular endothelium. When perfused with culture medium through the vascular channel, the BM Chip maintains human CD34+ cells and supports differentiation and maturation of multiple blood cell lineages over 1 month in culture. Moreover, it recapitulates human myeloerythroid injury responses to drugs and gamma radiation exposure, as well as key hematopoietic abnormalities found in patients with the genetic disorder, Shwachman-Diamond Syndrome (SDS). These data establish the BM Chip as a new human *in vitro* model with broad potential utility for studies of BM dysfunction.

---

The human BM is the site where all adult blood cells originate and thus BM dysfunction causes significant patient morbidity and mortality. It is a major target of drug‐ and radiation-related toxicities due to its high cell proliferation rates, and it is also affected by a variety of genetic disorders from congenital marrow failure syndromes to myeloid malignancies. While these abnormalities can be diagnosed and managed by monitoring peripheral blood counts, it is the proliferation and differentiation of hematopoietic cells in the marrow that is directly targeted in these disease states. Aside from invasive biopsies, there are no methods to study these responses *in situ* over time in human patients.

Various *in vitro* culture methods for human hematopoietic cells have been described, including culturing CD34+ hematopoietic progenitors in suspension (including methylcellulose-based assays)^1,2^, on stromal cell monolayers (e.g. Dexter culture and assays to assess long-term culture-initiating cells and cobblestone area-forming cells)^3,4^, or in perfused devices that may include bone or artificial scaffolds seeded with various stromal cells^5-9^. These *in vitro* methods, along with an extensive body of work in animal models, have been used to gain fundamental insight into the biology of hematopoiesis and hematopoietic stem cells^1,2,10^. However, the translational application of human *in vitro* culture methods to modeling hematopoietic injury and disease has been limited. Methylcellulose-based colony forming assays remain the workhorse, especially in the pharmaceutical industry, despite inherent limitations in the ability to manipulate the culture (e.g. add or remove drug) after the initial setup^3,11^. Other methods have largely focused on expanding CD34+ progenitors or differentiating specific hematopoietic lineages, which has led to interesting potential uses in cell therapy but their wider applicability is limited^6,12–16^. Thus, using existing *in vitro* BM models, it has not been possible to recapitulate a broad spectrum of clinically relevant responses of living marrow to drugs, radiation, and disease-causing mutations, which would greatly expand their applications to human health.

## Results

### Human BM Chip supports hematopoietic cell growth and CD34+ progenitor survival

The hematopoietic compartment of the marrow contains stem and progenitor cells that proliferate and differentiate into mature white and red blood cells through interactions with a variety of surrounding stromal cells (including BMSCs)^10,17^ and it interfaces directly with an endothelium-lined vasculature (**Fig. 1a**). The vascular compartment also contributes to BM function by producing angiocrine factors, supplying nutrients, and removing waste^10,18^. To model this organ functional unit, we adapted a previously described 2-channel, microfluidic Organ-ona-Chip (Organ Chip) device^19,20^ (**Supplementary Fig. 1a**). The top ‘hematopoietic’ channel was filled with a three-dimensional co-culture of human CD34+ cells and BMSCs in a fibrin gel while the bottom ‘vascular’ channel was lined by human umbilical vein endothelial cells (HUVECs) (**Fig. 1a**). The BM Chip was fed exclusively via medium perfusion through the vascular channel, analogous to the *in vivo* situation.

**Fig. 1.**
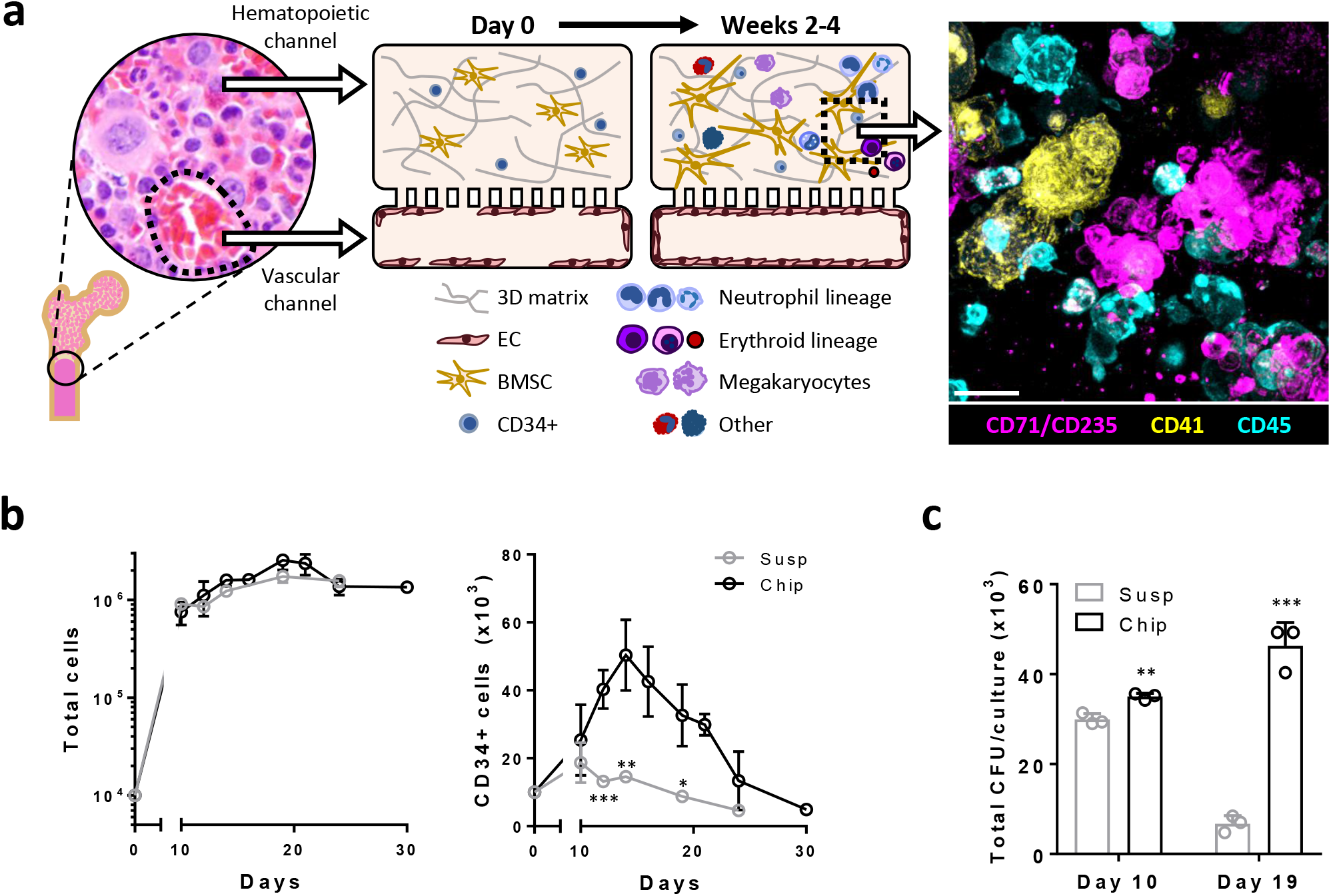
Primary human BM Chip supports hematopoietic cell growth and CD34+ progenitor survival over one month in culture. **a**, Schematic of human bone with a micrograph showing normal human BM histology (left) and a schematic cross-sectional view of the human BM Chip on day 0 after seeding showing singly dispersed CD34+ progenitors and BMSCs in a gel in the top channel with an incomplete vascular lining in the bottom channel (left middle). Within 2 weeks of culture, endothelial cells grow to cover all four sides of the lower channel and create a vascular lumen while CD34+ cells undergo expansion and multilineage differentiation (right middle), as illustrated by immunofluorescence image of a vertical cross section through the gel in the upper channel of the BM chip taken at day 14 (magenta: erythroid lineage; yellow: megakaryocyte lineage; blue: neutrophil and other hematopoietic lineages; Scale bar, 20 μm). **b**, Numbers of total cells (**left**) and CD34+ cells (**right**) measured over time in the BM Chip or standard 96 well plate suspension cultures using flow cytometry (n=3-17 chips or 3-6 wells per timepoint; data pooled from 5 independent experiments). **c**, Total colony forming units (CFUs) per culture when 1 × 10^4^ CD34+ cells were seeded into BM Chips versus standard 96 well plate suspension cultures. After 10 or 19 days, cells were harvested, plated in methylcellulose cultures, and total CFUs per BM Chip or well were quantified (n=3 per timepoint; day 10 and day 19 data from independent experiments). All data are presented as mean ± s.d. (*P<0.05; **P<0.01; ***P<0.001).

Over multiple weeks of culture, the CD34+ progenitors proliferated and coalesced to form a dense cellular microenvironment composed of multiple hematopoietic lineages, analogous to that observed in living BM (**Fig. 1a, Supplementary Fig. 1b**). The total number of cells present within the BM chip increased over 100‐ to 200-fold within 14 days of culture, after which cell numbers remained steady for 2 additional weeks (**Fig. 1b**). The number of CD34+ cells also increased ~5-fold over the first 2 weeks but then decreased in number as multilineage differentiation increased at later times (**Fig. 1b**). Compared to BM Chip cultures, CD34+ cell suspension cultures supported the growth of similar numbers of total cells but contained fewer CD34+ cells and displayed decreased progenitor function over time, as measured by methylcellulose colony forming ability (**Fig. 1b,c, Supplementary Fig. 1c**). Static fibrin gel co-cultures of the same CD34+ progenitors and bone marrow stromal cells in BM Chips also contained fewer CD34+ cells over time, including immunophenotypically immature CD38-CD34+ cells (**Supplementary Fig. 1d**).

### Continuous myeloid and erythroid cell proliferation and differentiation in the BM Chip

Flow cytometric analysis and Wright-Giemsa staining showed that cells within the BM Chip were predominantly of the neutrophil and erythroid lineages, and they displayed the full spectrum of maturation states (**Fig. 2a,b, Supplementary Fig. 1e,f**). Other blood cell types were also produced, including megakaryocytes and macrophages, although they were present in fewer numbers (**Supplementary Fig. 1e,f**). Lymphoid-specific growth factors were not included in the culture medium and, as expected, no lymphoid differentiation was observed (data not shown).

**Fig. 2.**
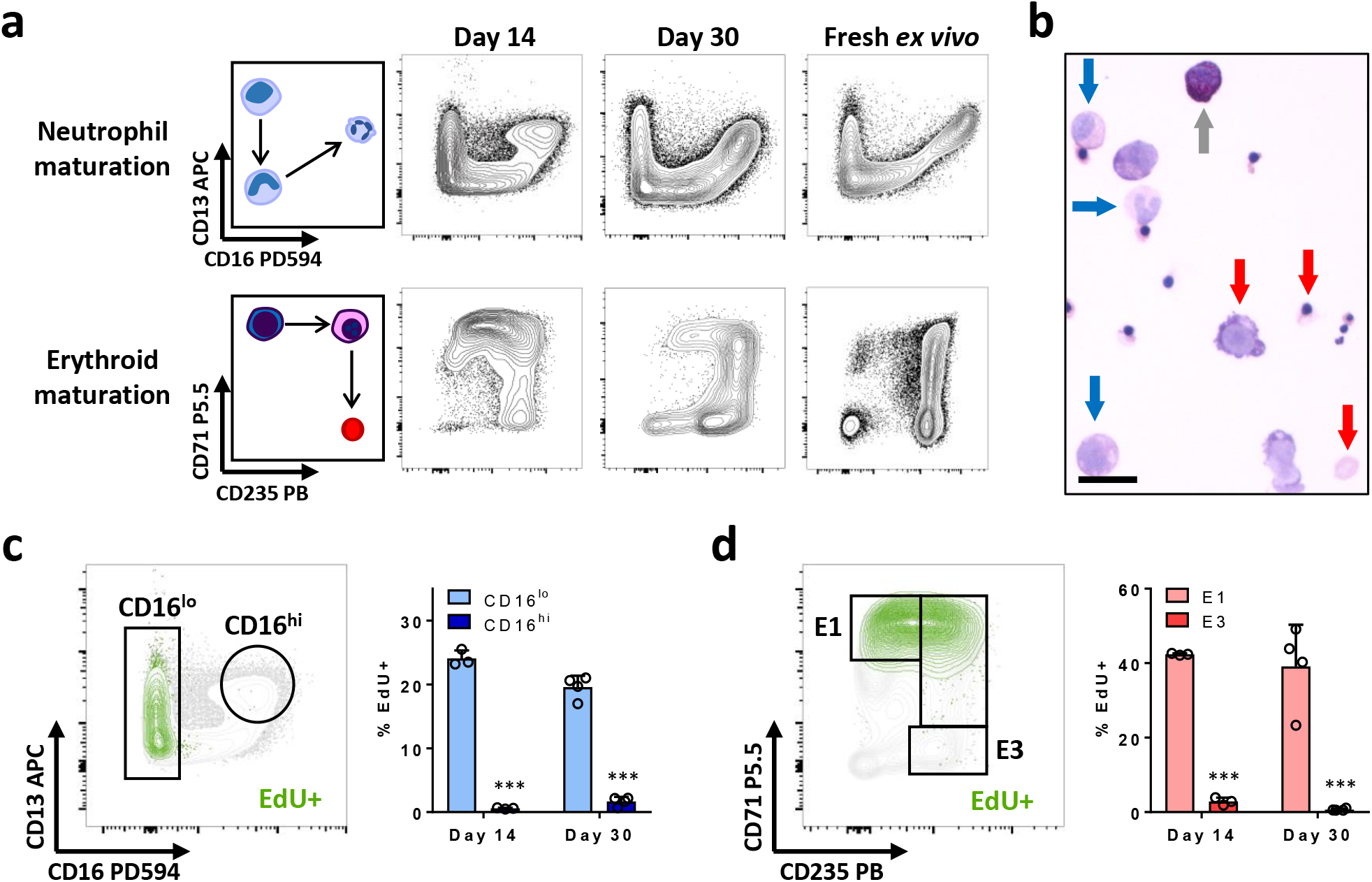
Continuous myeloerythroid proliferation and differentiation in the BM Chip. **a**, Schematic diagrams (left) and representative flow cytometry plots of neutrophil and erythroid maturation (right) showing similar phenotypic profiles between cells extracted from the BM Chip and from fresh human bone marrow. **b**, Wright-Giemsa stain of cells from a BM Chip at day 21 showing multiple cell types at varying stages of maturation (blue arrows, neutrophil lineage; red arrows, erythroid lineage; gray arrow, non-neutrophil granulocyte; Scale bar, 20 μm). **c**, Cell proliferation within the neutrophil lineage was assessed by a 2-hour EdU pulse immediately prior to cell harvesting. Flow plot illustrates representative gating strategy for immature CD16^lo^ and mature CD16^hi^ neutrophil subpopulations while EdU+ neutrophil lineage cells are highlighted in green. The percentage of EdU+ cells within the immature and mature neutrophil populations was quantified by flow cytometry at days 14 and 30. **d**, Cell proliferation within the erythroid lineage was similarly assessed in the same BM Chips. Flow plot illustrates representative gating strategy for immature CD71+CD235‐ (E1) and mature CD71-CD235+ (E3) erythroid subpopulations while EdU+ erythroid lineage cells are highlighted in green. The percentage of EdU+ cells within the immature and mature erythroid populations was quantified by flow cytometry at days 14 and 30. (n=3-4 chips per timepoint; data pooled from 2 independent experiments).

To better assess the dynamics of on-chip hematopoiesis, cells were labeled with a 2 hour pulse of EdU just before they were harvested and analyzed at days 14 and 30. EdU+ cells within the neutrophil lineage appropriately resided largely within the immature CD16^lo^ population as opposed to the mature CD16^hi^ population (**Fig. 2c**). Similarly, EdU+ cells of the erythroid lineage primarily exhibited an immature CD71+CD235‐ (E1) phenotype as opposed to a mature CD71-CD235+ phenotype (E3) (**Fig. 2d**). Importantly, EdU analysis at days 14 and 30 showed that the immature CD16^lo^ neutrophil and E1 erythroid cells maintained their proliferative nature over time (**Fig. 2c,d**). Furthermore, the proliferative fraction among total cells decreased as the BM Chip evolved to contain an increasing percentage of mature post-mitotic cells (**Supplementary Fig. 1g**) but the CD34+ population retained significantly higher EdU labeling, indicating continued CD34+ progenitor cell proliferation (**Supplementary Fig. 1g**).

In the course of these studies, we observed that a small fraction of cells spontaneously migrated or “intravasated” into the vascular channel over the first 2 weeks of culture, and these cells were markedly biased towards the myeloid lineage relative to the erythroid lineage when compared to cells remaining in the hematopoietic channel (**Supplementary Fig. 2a**). These intravasating cells also contained a 5-fold higher proportion of mature CD16^hi^ neutrophils (**Supplementary Fig. 2a**), which is reminiscent of the selective migration from marrow into peripheral blood by mature, but not immature, granulocytes that occurs *in vivo.* Importantly, plate-based cultures of CD34+ cells cannot mimic these effects and they also have known limitations with respect to diffusion-limited exchange of oxygen and nutrients, as well as removal of waste and soluble inhibitory factors^21^. In contrast, when we fabricated BM Chips using microfluidic devices that contained embedded oxygen sensors^22^, we found that oxygen levels remained constant at ~ 75% of atmospheric levels over 3 weeks in the BM Chip, whereas oxygen levels in static fibrin gel co-cultures containing the same cells dropped precipitously to near anoxic levels within the first week (**Supplementary Fig. 2b**). These findings confirm that the microfluidic BM Chip significantly improves oxygen delivery (and likely nutrient and waste exchange as well) compared to static plate-based cultures.

### BM Chip predicts lineage-specific drug toxicities and marrow recovery upon withdrawal

BM toxicity is a significant dose-limiting side effect for many drugs. Hence, there is a great need for human *in vitro* models that can predict drug-induced myelosuppression at patient-relevant exposures and model the dynamics of marrow recovery from injury. To explore the utility of the BM Chip as a model of drug toxicity, we tested AZD2811, which is a selective small molecule inhibitor of Aurora B kinase that is currently in Phase I clinical development in an encapsulated polymer nanoparticle form. Interestingly, when AZD2811 was delivered by intravenous infusion as the rapidly converting phosphate prodrug barasertib, it was found to produce different myelotoxicity profiles when the same total dose was administered over either 2 or 48 hours in Phase I clinical trials for various malignancies^23,24^. To mimic the infusion schedules used in those studies^23,24^, we infused the vascular channels of BM Chips beginning after 10 days of culture with clinically relevant amounts of AZD2811 over 2 or 48 hours (total Area Under the Curve [AUC] of 0.5, 1, or 2 μM.h in both cases). These doses were calculated to deliver a range drug exposures that reach those observed in the plasma of human patients who exhibited cytopenias^24^ (see **Methods** for details). The attainment of patient-relevant dosing in terms of AUC and overall shape was confirmed by comparing direct measurements of AZD2811 concentrations in BM Chip outlet medium, pharmacokinetic (PK) models of BM Chip exposures based on those measurements, and the simulated free AZD2811 concentration profiles in the plasma of an average patient (based on its known PK properties) at a range of clinical doses^24^ (**Fig. 3a, Supplementary Fig. 3**).

**Fig. 3.**
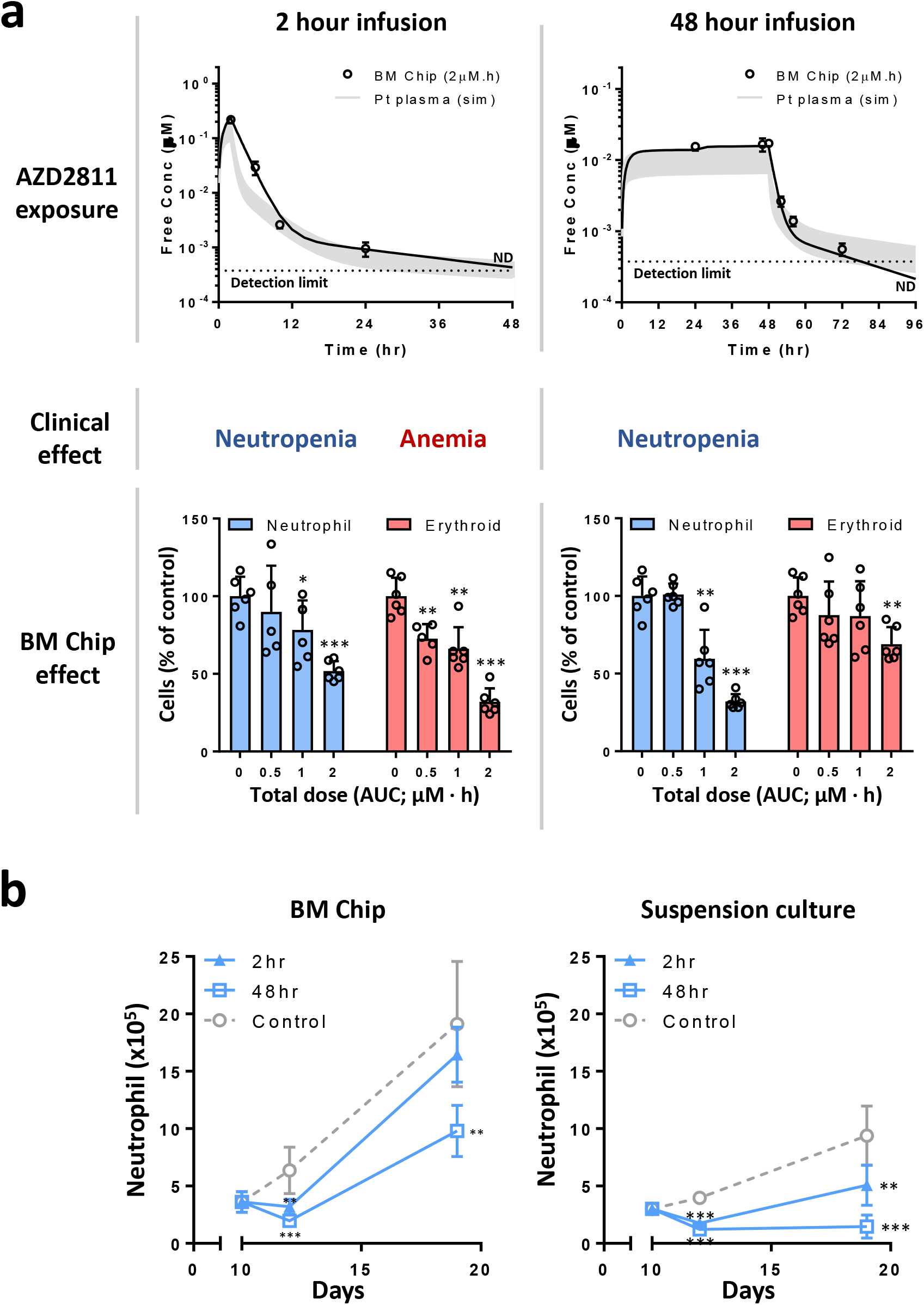
BM Chip predicts lineage-specific side effects of AZD2811 and models marrow recovery upon drug withdrawal. **a**, Top) Plasma levels of AZD2811 in an average patient simulated at a range of clinical doses^24^ for 2 hour (left) and 48 hour (right) infusions based on the known PK characteristics of AZD2811 (Pt plasma) compared with predictions of a PK model of BM Chip drug exposure (solid line) derived from experimentally measured drug concentrations (circles). The BM Chips were matured for 10 days, treated with 2-hour or 48-hour infusions of AZD2811, and then cultured again in drug-free medium. AZD2811 concentrations were measured by mass spectrometry in BM Chip outlet medium collected at various timepoints during and after drug infusion (data from the AUC = 2μM.h condition is shown here). Drug levels were below the detection limit (ND = not detectable) at 48 and 96 hours after the start of infusion for the 2-hour and 48-hour regimens, respectively. Dotted line represents the detection limit of mass spectrometry during these experiments. Middle) Hematologic side effects (neutropenia or anemia) previously reported in patients receiving 2-hour (left) or 48-hour (right) infusions of similar total doses of barasertib^23,24^ in Phase I clinical studies. Bottom) Graphs showing the effects of exposing the BM Chips on day 10 of culture to varying doses of AZD2811 for 2 hours (left) versus 48 hours (right) on total neutrophil (blue) and erythroid (red) cell numbers on day 12. Neutrophil and erythroid lineage cells were quantified by flow cytometry (n=6 chips per condition; data pooled from 2 independent experiments). **b**, Effects on neutrophil numbers in BM Chips (left) or suspension cultures (right) when treated at day 10 with the highest total dose of AZD2811 (2μM.h) over 2 or 48 hours and subsequently allowed to recover in drug-free medium. Neutrophil numbers were quantified by flow cytometry (n=6 chips/wells per condition at each timepoint; data pooled from 2 independent experiments; *P<0.05; **P<0.01; ***P<0.001).

Importantly, flow cytometric analysis revealed that the 2-hour and 48-hour infusions had different effects in the BM Chip. Dose-dependent toxicity in the erythroid lineage was greater following the 2-hour infusion regimen (**Fig. 3a**), which closely aligns with clinical data where anemia (CTCAE grade < 3) was only reported as an adverse event in patients receiving the 2hour infusion of barasertib^23,24^. In contrast, dose-dependent toxicity of the neutrophil lineage in the BM Chip appeared to be greater with the 48-hour infusion (**Fig. 3a**) than the 2-hour infusion (68% vs. 48% decrease at the highest tested dose; p < 0.001). This is also consistent with published clinical data showing a higher incidence of severe neutropenia (CTCAE grade ≥ 3) with drug infusions over 48 hours versus 2 hours at a given AUC^23,24^. Importantly, suspension cultures of the same CD34+ cells failed to reproduce the regimen-dependent erythroid toxicity (anemia) profile that was mimicked by the human BM Chip (**Supplementary Fig. 4a**).

Clinical trial data show that the hematologic side effects of barasertib were readily managed by withholding drug or stimulating with growth factors^23,24^, indicating that efficient marrow recovery occurs when administration of AZD2811 is ceased. Similarly, when we cultured the human BM Chip in drug-free medium for 7 additional days after administering the highest dose of AZD2811 (2 μM.h), we observed robust neutrophil recovery with both the 2 and 48-hour dosing regimens as well as erythroid recovery with the 2-hour regimen (**Fig. 3b, Supplementary Fig. 4b**). In contrast, this level of neutrophil recovery was not seen in plate-based CD34+ suspension cultures (**Fig. 3b**). These results suggest that the BM Chip could be used to model a full cycle of BM injury and recovery during preclinical drug evaluation and development, which up to now has required animal studies.

### Effects of drugs and ionizing radiation on mature versus immature hematopoietic cells

Aurora B kinase, the pharmacological target for AZD2811, is necessary for the proper alignment and segregation of chromosomes during mitosis as well as completion of cytokinesis^25^. Thus, inhibition of this kinase by AZD2811 should preferentially damage immature dividing precursors while sparing mature post-mitotic cells. Aside from doing a marrow biopsy, it would be difficult to assess this response in patients; however, we are able to test this directly in the human BM Chip. Indeed, the effect of AZD2811 was significantly greater on immature CD16^lo^ neutrophil precursor cells than on mature CD16^hi^ neutrophils with both dosing regimens (**Fig. 4a**). Furthermore, the anemia-inducing 2-hour infusion regimen caused preferential toxicity in early E1 erythroid progenitors versus more mature E3 erythroid cells (**Fig. 4b**).

**Fig. 4.**
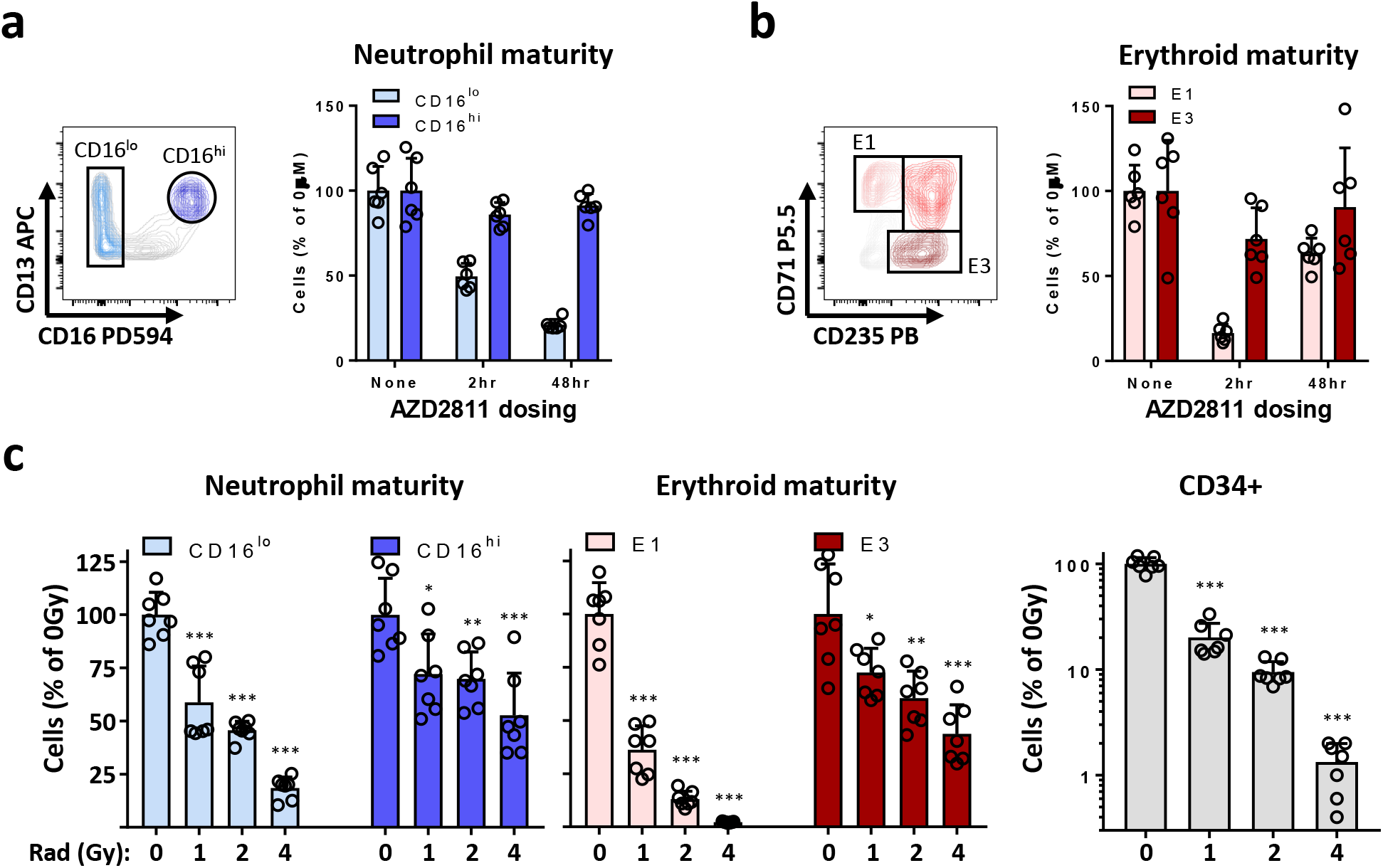
BM Chip enables assessment of AZD2811 and ionizing radiation-induced toxicity in hematopoietic cells at different stages of maturation. **a**, Representative flow cytometry plot (left) and graph (right) showing the preferential decrease in immature CD16^lo^ neutrophil lineage cells compared to mature CD16^hi^ neutrophil cells when BM Chips were treated on day 10 with the highest dose of AZD2811 (2μM.h) over 2 or 48 hours and analyzed on day 12 (n=6 chips per condition; data pooled from 2 independent experiments). **b**, Representative flow cytometry plot (left) and graph (right) demonstrating preferential toxicity to immature CD71+CD235‐ (E1) cells compared to more mature CD71-CD235+ (E3) erythroid cells by the 2-hour exposure to AZD2811 in the same BM Chips (n=6 chips per condition; data pooled from 2 independent experiments). **c**, Graphs showing neutrophil, erythroid, and CD34+ cell numbers determined using flow cytometry in BM Chips that were matured for 10 days, exposed to varying doses of ionizing radiation (0,1, 2, or 4 Gy), and then sacrificed at day 14. Note that radiation exposure preferentially reduced the numbers of immature CD16^lo^ neutrophil (left) and immature (E1) erythroid cell subpopulation (middle) compared to the more differentiated CD16^hi^ and E3 populations (n=7 chips; data pooled from 2 independent experiments; *P<0.05; **P<0.01; ***P<0.001).

The maturation-dependent effects we observed in response to drug exposure also would be predicted to occur in response to other types of myelotoxic stress, such as ionizing gamma (*γ*)-radiation. To test this, we exposed BM Chips to ionizing radiation doses ranging from 1 Gray (Gy), for which total body irradiation in humans results in mild recoverable cytopenias, to 4 Gy which causes 50% lethality over 60 days in the absence of supportive care^26^. BM Chips exposed to these levels of *γ*-radiation showed mildly decreased cell numbers at 1 Gy and severe toxicity at 4 Gy, matching human radiation sensitivities, and also displayed the predicted preferential toxicity of immature precursors as assessed by neutrophil, erythroid, and CD34+ cell numbers (**Fig. 4c, Supplementary Fig. 4c**). The responses of the BM Chip to AZD2811 and radiation-induced injury demonstrate that it can accurately predict nuanced hematopoietic toxicities with similar sensitivities seen in human patients caused by marrow stressors targeting disparate molecular pathways.

### Modeling Shwachman-Diamond Syndrome in the human BM Chip

Shwachman-Diamond Syndrome (SDS) is a genetic bone marrow failure syndrome which may also affect other organ systems, particularly the exocrine pancreas^27,28^. The majority of patients harbor biallelic mutations in the *SBDS* gene. The bone marrow is typically hypocellular for age and neutropenia (found in almost all patients) is the most common hematologic abnormality^29^. Hematologic dysfunction may involve other lineages (e.g. thrombocytopenia or anemia) and there is a predisposition to developing myeloid malignancies as well^27,28^. The study of hematopoiesis in the context of SDS has been limited because animal models do not faithfully recapitulate the human disease^30–34^. To assess the ability of the BM Chip to phenocopy the hematopoietic defects of SDS, we cultured CD34+ cells from two SDS patients (**Supplementary Fig. 5a**) together with normal BMSCs and endothelial cells in the 2-channel microfluidic device.

After 2 weeks of culture, SDS BM Chips displayed both broad defects in hematopoiesis and cell-type specific responses. Overall, significantly fewer cells developed in the SDS BM Chip compared to control BM Chips (**Fig. 5a**, **Supplementary Fig. 5b**), analogous to the hypoplastic phenotype often observed in SDS patient marrow biopsies^27,29^. This decrease in cell numbers was present in both the neutrophil and erythroid lineages (**Fig. 5b**). Maintenance of CD34+ cell numbers in the SDS BM Chip was also impaired (**Fig. 5c**), consistent with an intrinsic defect in CD34+ progenitor function in SDS patients, which reinforces existing literature that has demonstrated their impaired colony forming ability^35,36^. We noticed that neutrophils in the SDS BM Chip displayed an aberrant maturation pattern (**Fig. 5d**, **Supplementary Fig. 5c**). In contrast, no maturation defect was observed in the erythroid lineage, evidenced by similar numbers of mature (E3) erythroid cells in SDS and control BM Chips (**Fig. 5e**). Instead, there was an unexpected loss of early (E1) erythroid progenitors (**Fig. 5e**), and this was consistent between the two different SDS patients. Further study is needed to determine the generality and implications of these findings but these data demonstrate that SDS BM Chips display multiple hematopoietic abnormalities that parallel the clinical manifestations of neutropenia, anemia, and bone marrow failure in this disease.

**Fig. 5.**
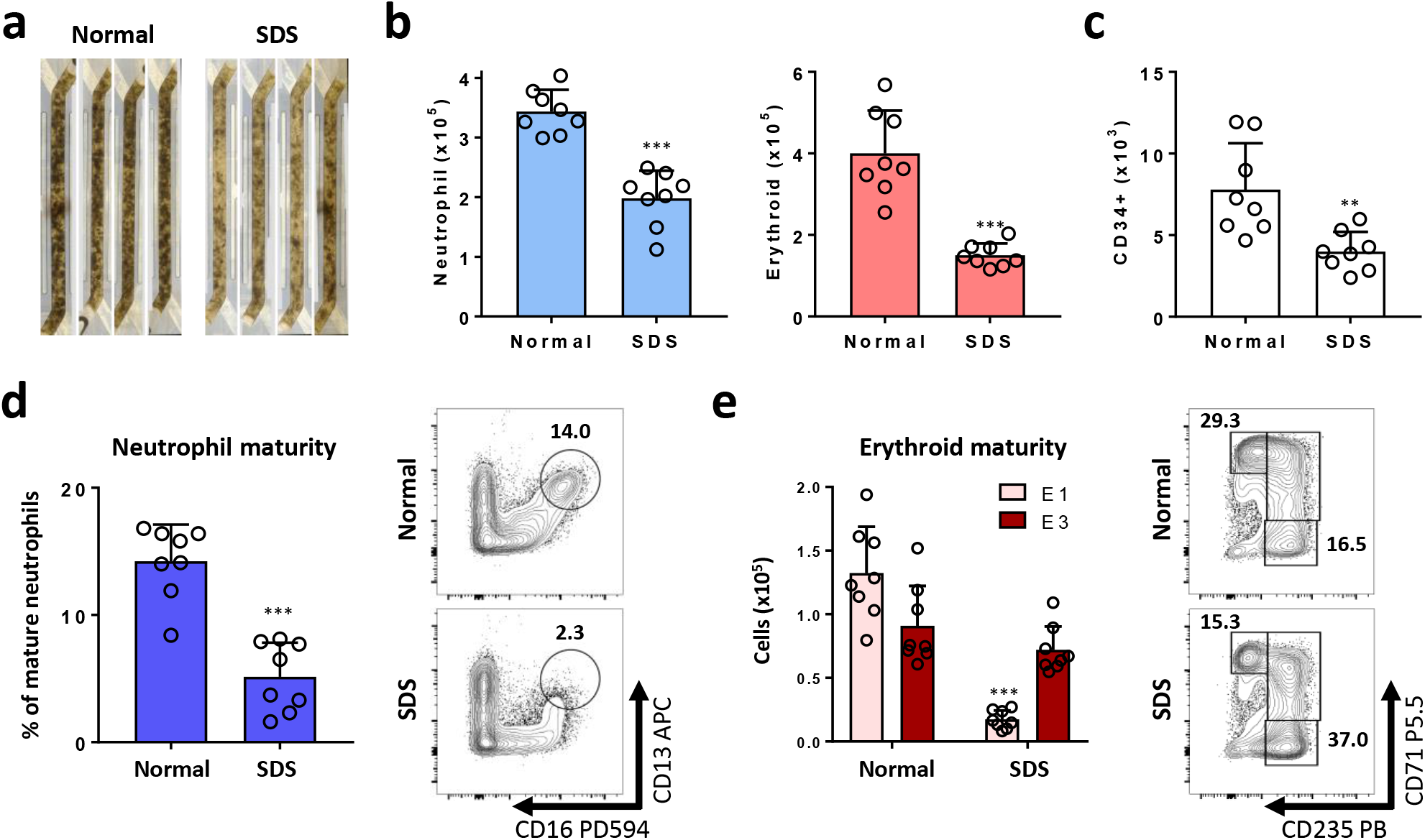
Modeling Shwachman-Diamond Syndrome in the human BM Chip. **a**, Photographs of BM chips seeded with CD34+ cells from normal controls versus SDS patients at 2 weeks of culture (n=4 chips per condition; representative of 2 independent experiments). Total neutrophil (**b**), erythroid (**b**), and CD34+ (**c**) cell numbers were quantified by flow cytometry. **d**, Graph (left) and representative flow plots (right) showing percentages of neutrophils with a mature CD16^hi^ surface phenotype in control versus SDS BM Chips, as quantified by flow cytometry. **e**, Graph showing the number of erythroid cells at different maturation states (left) and representative flow plots (right) depicting the percentages of the erythroid subpopulations (E1: immature, E3: mature), as quantified by flow cytometry. (**b-e**, n=8 chips; pooled from 2 independent experiments using cells from different SDS patients with 4 chips per experiment; *P<0.05; **P<0.01; ***P<0.001).

## Discussion

Taken together, our data show the microfluidic human BM Chip models key aspects of human hematopoiesis by supporting erythroid differentiation as well as myeloid development and mobilization over 1 month of culture while improving maintenance of CD34+ progenitors over traditional culture methods. The BM Chip was able to recapitulate the lineage-specific and maturation-dependent toxicity responses of human marrow to AZD2811 and ionizing radiation at human-relevant exposures. Finally, BM Chips seeded with SDS patient-derived CD34+ cells demonstrated hematopoietic dysfunctions that parallel key hematologic abnormalities observed in patients with Shwachman Diamond Syndrome.

It is worth noting that this study does not address issues of hematopoietic stem cell (HSC) presence or maintenance. Methods to effectively support, expand, or even create HSCs *in vitro* are active areas of investigation and would represent a milestone in hematopoietic culture systems^1,2,10,37–39^. This work, however, was directed towards creating an enabling technology with near-term translational potential as an effective model of human marrow pathophysiology. As the primary function of the bone marrow *in vivo* is the production of blood cells, we sought to balance proliferation, multilineage differentiation, and continued progenitor survival. In doing so, we were able to achieve extended myeloerythroid proliferation and maturation while also improving bulk CD34+ cell maintenance.

Microfluidic culture offers multiple advantages over conventional static culture methods, including improved nutrient exchange and waste removal. We found that the BM Chip, unlike a static gel culture, was able to maintain stable oxygen levels over time. Oxygen tension has been shown to play an important role in hematopoietic cell biology^40^. And while the experiments in this study were performed under atmospheric oxygen conditions, the ability to control oxygen levels enables future studies that match experimental oxygen tensions with those found in *in vivo* microenvironments^41–43^.

Careful consideration was given to how to best model Shwachman Diamond Syndrome given the limited availability of primary patient-derived cells. Ultimately, BM Chips were set up using SDS CD34+ cells but normal BMSCs as opposed to using BMSCs from SDS patients. This limited potential variability and allowed a better comparison to BM Chips seeded with normal CD34+ cells and BMSCs. Data obtained with this approach demonstrated interesting cell-intrinsic defects of CD34+ progenitors from SDS patients, which are known to play a central role in SDS pathology because hematopoietic stem cell transplant cures the marrow phenotype in SDS^28^. However, future experiments could examine possible contributions of the stromal microenvironment, which is a subject of active investigation in the field^31,35,44^.

Relatively little is known about how *SBDS* mutation affects the differentiation of human hematopoietic cells in the marrow. The impaired neutrophil maturation we observed in the SDS BM Chips aligns with prior work showing impaired function of peripheral blood neutrophils in SDS patients^36,45,46^ and it is also supported by results from SDS inhibition and knockout studies in mice^32,33^. The decrease in erythroid precursors was unanticipated, given that neither patient had clinically documented anemia. Nevertheless, the finding was consistent between two patients with different genetic abnormalities. It is possible, in light of a prior study showing impaired erythroid development upon *SBDS* knockdown in normal and immortalized human cells^47^, that increased erythropoietic stress may be a feature of hematopoietic dysfunction in SDS. However, further study is needed to confirm the physiologic significance of these data.

In summary, the vascularized human BM Chip represents a new *in vitro* model of human hematopoiesis that recapitulates many clinically relevant features of bone marrow pathophysiology in response to drugs, ionizing radiation, and genetic mutation. Future use of the BM Chip may reduce the incidence of unexpected myelotoxicity in the drug discovery process and the use of chips containing patient-derived cells could advance personalized medicine for patients with genetic BM failure disorders. This technology could also prove valuable in the study of a broader variety of marrow pathologies for which there are no current effective models.

## Methods

### Chip fabrication and preparation

The device was fabricated using polydimethylsiloxane (PDMS; SYLGARD^®^ 184 silicone elastomer kit) with previously described soft lithography techniques^19,48^ or obtained from Emulate, Inc. The chip design, which was similar to previously published devices^19^, included apical and basal microchannels with dimensions of 1 × 1 × 16.7 mm and 1× 0.2 × 16.7 mm (w × h × l), respectively, that were separated by a 50 μm thick porous PDMS membrane (7μm diameter holes with 40μm spacing). Both channels were washed with 70% ethanol, filled with 0.5 mg/mL sulfo-SANPAH solution (Thermo Fisher Scientific, A35395) in 50mM HEPES pH 8 and placed under a UV lamp (Nailstar, NS-01-US) for 20 minutes to activate the surface.The microchannels were then rinsed sequentially with 50 mM HEPES buffer and cold PBS, and then filled with a coating solution of PBS containing 100 μg/ml fibronectin (Gibco, 33016-015) and 50 μg/ml collagen (Sigma, C2249) and placed at 37°C for 2 hours. The solution was then aspirated from the chip and it was allowed to air dry overnight.

### Organ Chip Culture

The apical channel was first seeded with 1×10^4^ primary human CD34+ cells and 5 × 10^3^ BMSCs in a solution containing 5 mg/ml fibrinogen (Millipore, 341578), 0.2 mg/ml collagen I (Sigma, C2249), 25 μg/ml aprotinin (Sigma, A3428), and 0.5 U/ml thrombin (Millipore, 605195), that was allowed to gel on-chip. To prevent premature crosslinking, medium containing fibrinogen and aprotinin was prepared separately from the medium containing cells, collagen, and thrombin. The two solutions were then carefully mixed at a ratio of 1:3 (vol:vol) to avoid bubbles, immediately pipetted into the apical channel (25uL per chip), and allowed to gel for 20-30 minutes at room temperature. HUVECs (Lonza, CC-2517 or C2519A) were seeded either at this point or at day 8 by adding medium containing the cells (2-3×10^6^/ml; 30uL/chip) into the lower fibronectin‐ and collagen-coated channel followed by immediate inversion of the device and static incubation at 37°C for 2 hours to promote adhesion to the central membrane. The devices were then placed right-side up and apical channel ports were sealed with sterile tape. The entire BM Chip was then fed via medium perfused only through the endothelium-lined channel using a 16-channel peristaltic pump (Ismatec, ISM 938D) with 0.25 mm inner diameter (ID) pump tubing (Ismatec, 95723-12) with 19G bent metal pins at 1.42-1.6μL/min (pump speed varied depending on manufacturer’s lot of tubing). Outflow medium was collected using 0.89 mm ID tubing (Cole-Parmer, 95809-26) with 19G bent metal pins into a sterile reservoir. All parts were sterilized using oxygen plasma (Diener ATTO; 0.3 mbar pressure, 50W power and 2 min plasma time). Pumps were set to intermittent flow and pushed at 200 μL/day for the first seven days and 400μL/day thereafter. For drug studies, the volume was increased to 1.6 mL/day when drug treatment was initiated.

In all studies, we used SFEM II medium (STEMCELL Technologies, 09655) supplemented with 10% FBS (Gibco, 10082-147), 100 U/l penicillin and 100 mg/ml streptomycin (Pen/Strep; Gibco, 15140-122), 12.5 μg/ml aprotinin (Sigma, A3428), 20 ng/ml EPO (PeproTech, 100-64), 1 ng/ml G-CSF (PeproTech, 300-23), 100 ng/ml Flt3-L (PeproTech, 300-19), 100 ng/ml TPO (PeproTech, 300-18), 50 ng/ml SCF (PeproTech, 300-07), and select EGM-2 BulletKit (Lonza, CC-4176) components (hFGF-B, VEGF, R3-IGF-1, hEGF, ascorbic acid, and heparin) according to the manufacturer’s instructions. Medium was degassed under vacuum for at least 15 minutes before use to minimize bubble formation. Chips were kept humidified with damp tissues (replaced every 2 days; dampened with sterile water containing Lysol No-Rinse Sanitizer diluted 500x) placed on top of the chips and the whole setup was covered with aluminum foil.

### Static gel and suspension cultures

For static gel cultures, the same numbers of CD34+ cells and BMSCs as used in the BM Chips were seeded into individual wells of a 96 well plate (Corning, 353072) in 50μL of the same fibrin gel in order to cover the entire well surface. After crosslinking, 200μL of medium was added on top of the gel; the medium was fully replaced on day 3 and then daily starting at day 5. For suspension cultures, the same numbers of CD34+ cells were seeded into individual wells of a 96 well plate in 250μL of medium, and 50% of the medium volume was replaced daily.

### Oxygen measurements

BM Chips with embedded oxygen sensors were created as recently described^22^. For static gel culture measurements, an oxygen sensor spot (PreSens Precision Sensing GmbH, SP-PSt3-NAU) was placed at the bottom of a 96 well flat bottom plate before cell seeding. Cells were then cultured as described above. Oxygen measurements were optically measured daily using an OXY-4 mini instrument (PreSens Precision Sensing GmbH).

### Hematopoietic CD34+ cell isolation

Mobilized peripheral blood and leukapheresis product were anonymously collected from donors undergoing stem cell mobilization at the Massachusetts General Hospital (MGH) under Institutional Review Board approved protocol #2015P001859. Mononuclear cells were purified via Histopaque 1077 gradient (Sigma, 10771). CD34+ cells were isolated using positive magnetic bead selection using CD34 MicroBead Kit (Miltenyi Biotec, 130-046-702) and LS columns (Miltenyi Biotec, 130-042-401) according to manufacturer recommendations. CD34+ purity routinely exceeded 85% as assessed by flow cytometry. Aliquots of 3-5 × 10^5^ cells were frozen in RPMI 1640 (Gibco, 12633-012) + 10% DMSO (Sigma, 41640) + 10% FBS (Gibco, 10082-147) using the CoolCell XL (Corning) at −80°C and then transferred to liquid nitrogen cryogenic storage (VWR, CryoPro). Upon thawing, CD34+ cell viability was >90% as assessed by trypan blue (Lonza, 17-942E) and the yield was typically ~60% of frozen cells.

### Flow cytometry analysis

Cells were harvested from BM Chips and static gel cultures after digestion using 2.5 mg/mL nattokinase (Japan Bio Science Laboratory, NSK-SD), 1 mg/mL collagenase type I (Gibco, 17100-017), 25mM HEPES (Thermo Fisher Scientific, 15630-080), and 10% FBS in DMEM (Gibco, 11885-084). To digest BM Chips, one port of the bottom channel was plugged with a 200μL filter tip and 100μL of digestion solution was added via the other port. The tape on top of the apical channel ports was removed and a 200μL filter tip containing 50μL of digestion solution was then plugged in each apical channel port. After incubation for 1 hour at 37°C, the gel was broken up by pipetting the digestion solution in the chip up and down. If necessary, an additional 30-60 min of incubation was allowed. To digest static gel cultures, all medium was removed and 100μL of digestion solution was added on top; after incubation for 1 hour at 37°C, cells were harvested or allowed an additional 30 min of digestion if necessary.

Harvested cells were centrifuged, resuspended in flow staining buffer composed of 1% FBS (Gibco, 10082-147), 25mM HEPES (Thermo Fisher Scientific, 15630-080), 1mM EDTA (Thermo Fisher Scientific, 15575-020), and 0.05% sodium azide (VWR, BDH7465-2) in DPBS (Gibco, 14190-144), and filtered through 105 μm pore nylon mesh (Component Supply Co, U-CMN-105-A). Antibody staining was performed for 30 min in 96 well V-bottom plates (Nunc, 249944) in 100μL of staining buffer using one of 2 panels:

Panel 1: anti-CD235a-Pacific Blue (HI264 clone, BioLegend, 349108, dilution 1:80), anti-CD15-Brilliant Violet 510 (W6D3 clone, BioLegend, 323028, dilution 1:50), anti-CD45-Brilliant Violet 570 (HI30 clone, BioLegend, 304034, dilution 1:50), anti-CD33-PE (WM53 clone, BioLegend, 303404, dilution 1:50), anti-CD16-PE/Dazzle 594 (3G8 clone, BioLegend, 302054, dilution 1:80), anti-CD41-PE/Cy5 (HIP8 clone, BioLegend, 303708, dilution 1:133), anti-CD71PerCP/Cy5.5 (CY1G4 clone, BioLegend, 334114, dilution 1:50), anti-CD34-PE/Cy7 (581 clone, BioLegend, 343516, dilution 1:50), anti-CD13-APC (WM15 clone, BioLegend, 301706, dilution 1:80).

Panel 2: anti-CD45-Brilliant Violet 570 (HI30 clone, BioLegend, 304034, dilution 1:50), anti-CD34-PE/Cy7 (581 clone, BioLegend, 343516, dilution 1:50), anti-CD38-APC (HB-7 clone, BioLegend, 356606, dilution 1:80).

Both panels included 20 nM Syto16 (Thermo Fisher Scientific, S7578, dilution: 1:500), Zombie NIR dye (BioLegend, 423106, dilution: 1:500), Fc Block (BioLegend, 422302, dilution 1:20), Monocyte block (BioLegend, 426103, dilution 1:20), and Brilliant stain buffer (BD, 566385, dilution 1:20). Additionally, 5 x 10^3^ counting beads (Spherotech, ACRFP-100-3) were added to each sample to enable quantification of cell numbers. Stained cells were analyzed using the LSRFortessa (BD Biosciences). Results were analyzed using FlowJo V10 software (Flowjo, LLC). A representative gating strategy is described in **Supplementary Fig. 6**.

### Immunofluorescence microscopy

BM Chips were fixed by plugging one bottom channel port with a 200μL filter tip and gently adding 200μL of 2% paraformaldehyde (Electron Microscopy Sciences, 15730), 25mM HEPES (Thermo Fisher Scientific, 15630-080) in DPBS (Gibco, 14190-144) via the other bottom channel port. After 2 hours, the fixation solution was removed and replaced with DPBS. The chips were then kept at 4°C until sectioning at 500μm with a vibratome (Leica, VT1000S). Sections were blocked/permeabilized using 0.1% Triton X-100 (Sigma, T8787), 5% Normal Goat Serum (Jackson Immunoresearch, 005-000-121), 25mM HEPES in DPBS for 30 minutes at room temperature. Samples were then stained overnight at 4°C in the same buffer containing Fc Block (BioLegend, 422302, 1:20), anti-CD41 Alexa Fluor 488 (Biolegend, 303724, 1:20), anti-CD71 purified (Biolegend, 334102, 1:400), anti-CD235 purified (Biolegend, 349102, 1:400), anti-CD45 Alexa Fluor 594 (Biolegend, 304060, 1:20). The next day, after 3 washes of PBS, sections were then stained overnight at 4°C in the same buffer containing goat anti-mouse IgG2a Alexa Fluor 555 (ThermoFisher, A-21137, 1:500), and Hoechst 33342 (Life Technologies, H3570, 1:10,000). Images were taken using a laser scanning confocal immunofluorescence microscopes (Zeiss TIRF/LSM 710) with a 405-nm diode laser, a 489–670 nm white light laser, 488 nm and 496 nm argon laser and coupled to a photomultiplier tube or HyD detectors. Acquired images were analyzed using IMARIS software (Bitplane).

### Cell growth assay

Growing cells were detected using the Click-iT™ Plus EdU Alexa Fluor™ 350 Flow Cytometry Assay Kit (Thermo Fisher Scientific, C10645). BM Chips were perfused with medium containing 10μM EdU for 2 hours immediately before cell harvesting as described above. After surface staining with panel 1 antibodies, samples were fixed with 100 μl of 2% paraformaldehyde (Electron Microscopy Sciences, 15730), 25mM HEPES (Thermo Fisher Scientific, 15630-080) in DPBS (Gibco, 14190-144) for 20 min at room temperature. Samples were then processed as per manufacturer’s protocol.

### Wright-Giemsa staining

1×10^5^ cells in 200 μl DPBS were spun (300 rpm, 5 minutes, RT) onto Superfrost Plus slides (VWR, 48311-601) using a Cytospin 4 (Thermo Fisher Scientific) cytocentrifuge and air dried. Slides were placed for 4 minutes in full-strength staining solution (Sigma, WG16) and then moved to diluted staining solution (1 part WG16 to 5 parts DPBS) for 30 minutes. Slides were then washed in deionized water, air dried, and cover slipped using Cytoseal 60 (Electron Microscopy Sciences, 18006). Representative images were taken using a Zeiss Axioplan 2 microscope with AmScope MU1830 digital camera.

### Colony forming unit (CFU) assay

CFU assays were performed using MethoCult^TM^ medium (STEMCELL Technologies, 04434) in meniscus-free 6 well plates (STEMCELL Technologies, 27370) as per manufacturer’s suggestions. Each well contained total cells sufficient to include 50-70 CD34+ cells as calculated from same-day flow cytometry results. Plates were placed into a secondary humidified chamber within an incubator and analyzed using the STEMvision instrument and software (STEMCELL Technologies) after 14 days.

### Drug treatment

AZD2811 (formerly known as AZD1152 hydroxy-quinazoline pyrazole anilide) was provided by AstraZeneca in DMSO and stored at −20°C. Fresh medium with drug at the indicated concentrations was prepared daily and added to BM Chip reservoirs and plate cultures for the indicated times. To stop drug treatment, BM Chips were flushed at maximum pump speed (using the MAX/CAL button) for 30 seconds while suspension cultures were harvested, diluted with additional medium, centrifuged at 300g for 10 min, and replated in fresh medium. Dosing was based on results from a Phase I clinical trial^24^, which showed that cytopenia-inducing doses of drug corresponded to a total Area Under the Curve (AUC_Total_) per dose of 4776 ‐ 13520 ng.h/mL in human plasma. Accounting for the molecular weight (507 g/mol) and protein binding of this drug (4% fraction unbound in human plasma; internal AstraZeneca testing), this corresponds to hematologic side effects being induced by free drug AUC_Total_ amounts of 0.376-1.06 μM.h. Correcting for protein binding of AZD2811 in the BM Chip medium (37.5% fraction unbound as determined by mass spectrometry) results in a corresponding AUC_Total_ range in BM Chip medium of 1.00 −2.83 μM.h. Thus, BM Chips were dosed at 0.5, 1, and 2 μM.h to match patient exposure levels.

### Mass spectrometry analysis of AZD2811

Drug concentrations in medium samples were determined by liquid chromatography-mass spectrometry/mass spectrometry (LC-MS/MS). Briefly, 50 μL of each sample was dispensed into a 96-well plate followed by addition of 250 μL of protein precipitation solution (80:20 acetonitrile:methanol) containing internal standard (clozapine). Plates were vortexed for 5 min and then centrifuged at 2150 × g for 5 min at 4 °C. Clear supernatant solutions (225 μL) were transferred to clean plates and dried to completeness under nitrogen, reconstituted in 80:20 10mM ammonium formate in water: acetonitrile (200 μL) and analyzed using LC-MS/MS. Liquid chromatography was performed using a Waters XBridge C18 3.5um, 30 × 3mm. The ionization was conducted in positive mode and the m/z transitions were 507.9/130.2 for AZD2811. Mass spectrometry was performed using an AB Sciex 6500 Triple quadrupole mass spectrometer (AB Sciex, Foster City, CA, USA) operated in electrospray ionization mode. Data were acquired and analyzed using Analyst software (v 1.6.2).

The unbound fraction of AZD2811 in culture medium was determined using equilibrium dialysis. Medium samples spiked with 1μM of compound were dialyzed using the Rapid Equilibrium Dialysis device (Thermo Fisher Scientific Inc, Rockford, IL) by transferring 300 μL of spiked medium to the dialysis unit’s medium chamber and 500 uL of PBS to the buffer chamber. The dialysis unit was incubated at 37°C and 400 rpm for 18 hours. At the end of the incubation, 25 μL of each sample from the medium and buffer chambers was transferred to a 96 well plate. Blank medium or buffer (25 μL) was added to the buffer or medium samples, respectively, to yield identical matrices. Samples were then quantified using LC-MS/MS, as described above, and the unbound drug fraction (F_unbound_) in the medium was calculated as follows: F_unbound_ = [Buffer chamber] / [Medium chamber].

### PK modeling and simulation

All model fitting and simulations were performed using WinNonlin Phoenix NLME 6.4A. Clinical pharmacokinetic profiles of AZD2811 plasma concentration versus time for a population average patient were simulated at each Phase 1 study dose level (75, 100 & 150 mg administered by 2 hour infusion; 100, 150 & 225 mg administered by 48 hour infusion) using the 3-compartment population PK model for AZD2811 described by Keizer et al^49^. The free fraction in human plasma was assumed to be 0.04 (AstraZeneca derived value).

For the BM Chip, a three-compartment PK model was fitted to the LC-MS/MS quantified AZD2811 concentrations. The central compartment in the model was viewed as representing the vascular channel and therefore the volume was fixed at the physical volume of the channel (20 μL) and the clearance fixed at the flow rate of medium through the channel (72 μL/hr). The first peripheral compartment was viewed as representing the hematopoietic channel and the volume fixed at the physical volume of the channel (25 μL). The rate and extent of distribution into the hematopoietic channel was described using a distribution clearance (Cl2) and a partition coefficient (Kphem) with both parameter value estimates being obtained during the model fitting process. Distribution into a second peripheral compartment (described by parameters V3 and Cl3) was found to be required to describe the second phase in the outlet medium concentration profile during the washout of drug from the system. Model parameter value estimates (K_phem_ = 4.5; Cl2 = 99 mL/hr; V3 = 163 μL; Cl3 = 5.6 μL/hr) were derived by fitting the PK model to the combined outlet medium concentration data from all AZD2811 dosed BM Chips using a naïve-pooled approach. Residual unexplained variability (between model and observed concentration data) was modelled using a multiplicative error model.

### Radiation exposure

For radiation exposure studies, BM Chips were disconnected from peristaltic pumps, transported in sterile containers to an irradiation facility at Boston Children’s Hospital, and exposed to a single selected *γ*-irradiation dose (Cs-137; Gammacell 40 Exactor) at 0.98 Gy/min. BM Chips were then reconnected to the peristaltic pumps, flushed for 5 seconds at maximum pump speed to remove bubbles, and placed in the incubator.

### Shwachman-Diamond Syndrome CD34+ cells

Written informed consent was provided by patients and normal donors in accordance with the Declaration of Helsinki. Primary human-derived bone marrow mononuclear cells were collected and frozen according to research protocols approved by the Institutional Review Board of Boston Children’s Hospital, Boston, MA. Upon thawing, SDS patient or normal donor-derived bone marrow mononuclear cells were allowed to recover in StemSpan SFEM II (STEMCELL Technologies, 09655) supplemented with recombinant human cytokines (all from PeproTech): 100ng/mL Flt3 (AF-300-19-50UG), 100ng/ml SCF (AF-300-07-50UG), 100ng/ml TPO (AF-300-18-50UG), 20ng/ml IL3 (AF-200-03-50UG). After 30-32 hours of recovery, CD34+ cells were sorted using magnetic beads as above and cultured in the BM Chip as described before.

### Statistical Analyses

All graphs depict mean ± standard deviation (s.d.) and tests for differences between two groups were performed using two-tailed unpaired Student’s t-test. Prism 7 (GraphPad Software) was used for statistical analysis.

### Data Availability

The authors declare that the data supporting the findings of this study are available within the article and its supplementary information files.

## Acknowledgments

This research was sponsored by funding from the US Food and Drug Administration grant HHSF223201310079C, Defense Advanced Research Projects Agency under Cooperative Agreement Number W911NF-12-2-0036, AstraZeneca, and the Wyss Institute for Biologically Inspired Engineering (to D.E.I.); US National Institutes of Health (R24 DK099808 and 5U01HL134812 to A.S., R01 DK102165 to C.D.N., and 5T32CA009216-37 training grant to D.B.C.); and the Department of Defense (W81XWH-14-1-0124 to C.D.N.). Additional funding was provided by the Dana-Farber Cancer Center Claudia Adams Barr Award (C.E.J.) and the EPSRC Centre for Innovative Manufacturing in Regenerative Medicine (A.R.). The authors would like to thank S. Sweeney for helpful discussions, and M. DeLelys, R. Matthews, J. Houston, J. Patel, D. Kingman, A. Shay, J. Graham, S. Chung, T. Spitzer, and F. Preffer at Massachusetts General Hospital for their invaluable help in working with patient samples.

## Author Contributions

D.B.C. and V.F. participated in the design and performance of all experiments and analysed the data, working with D.E.I, who also supervised all work. Y.M. helped design and perform experiments. R.D., P.P-D., D.F., A.M., and L.E. helped to design experiments relating to drug testing, performed mass spectrometry and PK modeling, analysed data, and helped write the manuscript. O.V.B. and C.E.J. helped design, perform, and interpret SDS-related studies with the input and supervision of A.S. and C.D.N.K.M. provided access to patient material for SDS-related studies. L.S.M.T. and A.R. helped conceive the BM Chip design and performed experiments. A.J. helped perform radiation-related studies. E.C., C.F.N, Y.C., and S.C. fabricated and participated in the design of the BM Chip with the input and supervision of R.N. and D.E.I. E.C., S.J-F., and S.C. helped perform oxygen studies. R.P.H. provided scientific supervision as well as access to patient material for all studies. O.L. and R.P-B. helped design experiments, interpret data, and supervised all work. D.B.C., V.F., and D.E.I. prepared the manuscript with input from all authors.

## Supplementary Figure Legends

**Supplementary Fig. 1.**
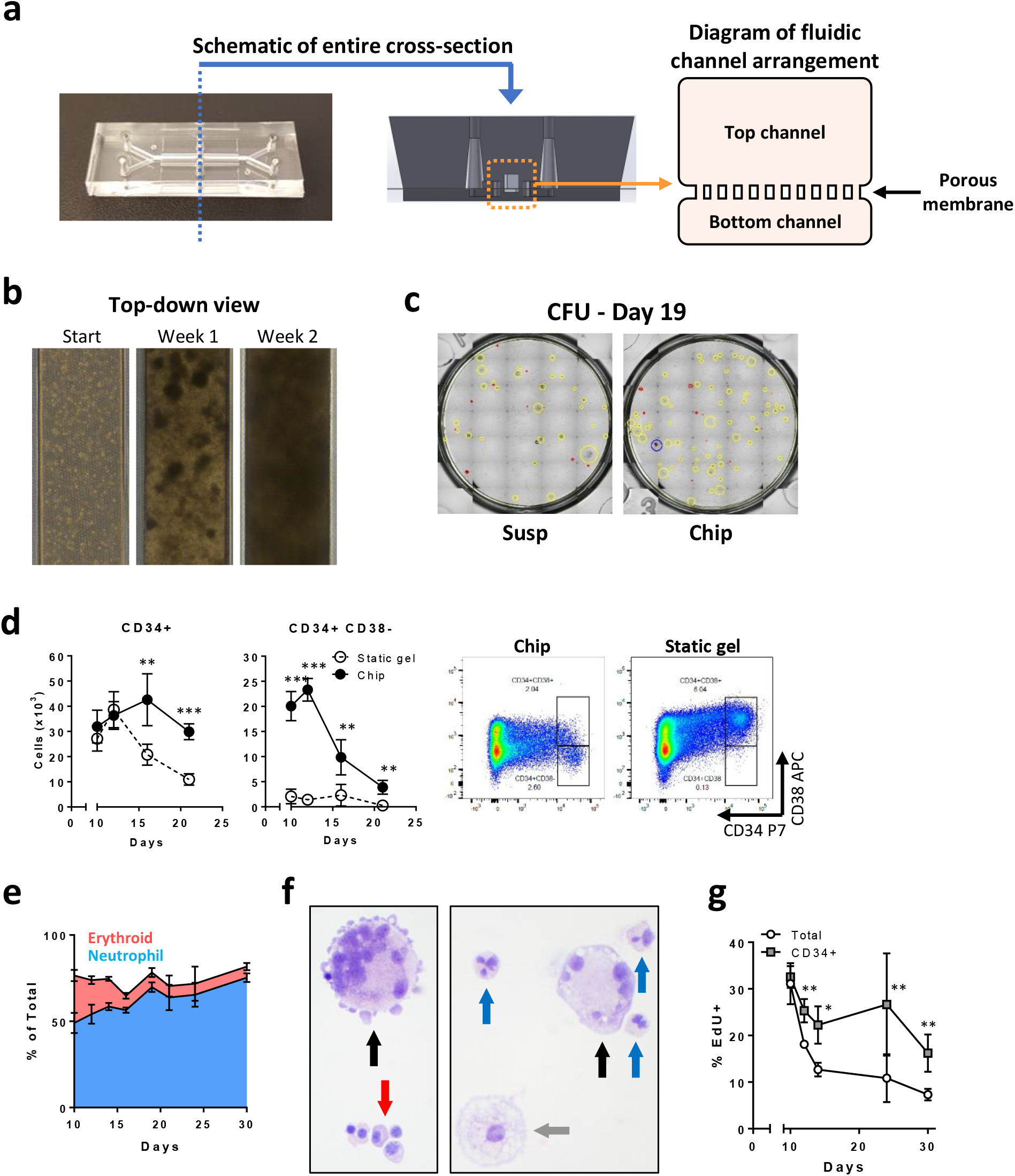
**a**, Photograph of an optically clear PDMS Organ Chip (left) used to create the human BM Chip along with a schematic of the vertical cross-section of the chip (middle) and a magnified diagram of the fluidic channels. **b**, Bright field micrographs of the top channel of the BM Chip viewed over time from above highlighting the development of cell clusters at week 1 that coalesce to form a dense cellular microenvironment by week 2. **c**, Representative photographs of methylcellulose cultures at 2 weeks that were derived from cells harvested at day 19 from suspension cultures versus BM Chips. **d**, Graphs (left) and representative flow cytometry plots at day 10 (right) showing total numbers of CD34+ and CD34+ CD38‐ progenitors in BM Chips or static gel cultures in a 96 well plate that were initially seeded with equivalent numbers of CD34+ cells and BMSCs (n=3-10 chips or wells per timepoint; data pooled from 4 independent experiments) **e**, Percentage of total cells represented by neutrophil (blue) versus erythroid (red) lineage cells in the BM Chip over time (n=3-17 chips per timepoint; pooled from 5 independent experiments). **f**, Wright-Giemsa stained cells isolated from BM Chips at day 14 illustrating multilineage differentiation (red arrow, group of maturing erythroid cells; blue arrows, mature segmented neutrophils; gray arrow, macrophage; black arrows, megakaryocytes). **g**, Cell proliferation among total and CD34+ cells in BM Chips assessed by a 2-hour EdU pulse immediately prior to cell harvesting at the indicated timepoints (n=3-9 chips per timepoint; data pooled from 3 independent experiments). (**d,g**: *P<0.05; **P<0.01; ***P<0.001).

**Supplementary Fig. 2.**
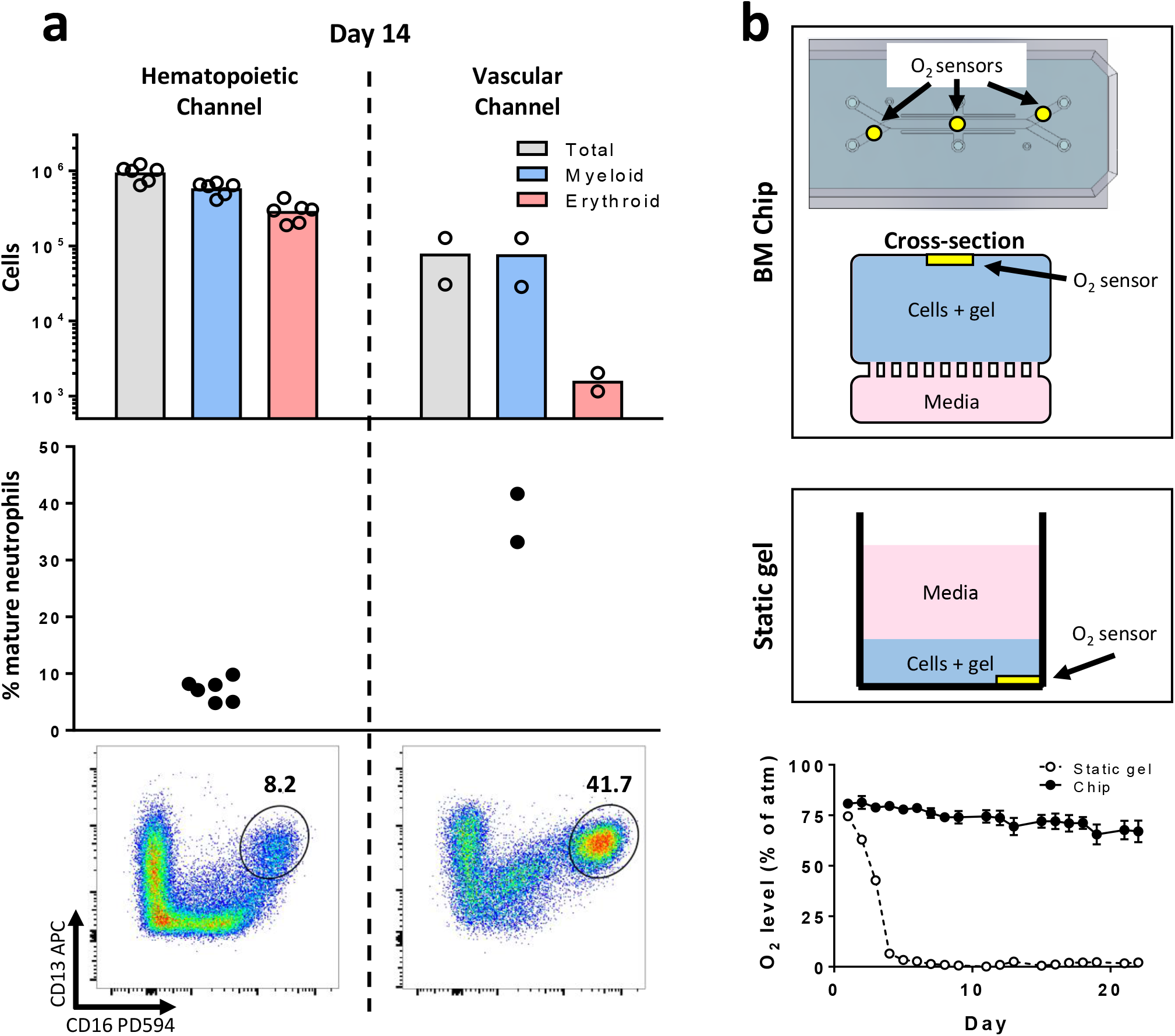
**a**, Graphs showing total, myeloid, and erythroid cell numbers (top) within individual hematopoietic channels or combined from 3-4 vascular channels of BM Chips measured by flow cytometry after 2 weeks of culture, as well as the percentage of neutrophils with a mature CD16^hi^ surface phenotype (middle) and representative flow plots (bottom) showing neutrophil maturation status (hematopoietic channel, n=6 chips; vascular channel, n=2 pooled from 3 or 4 chips each; data pooled from 2 independent experiments). **b**, Schematic showing location of oxygen sensors in the modified BM Chip (top) and 96 well static gel culture (middle). Graph at bottom shows oxygen levels that were optically measured in the two cultures on a daily basis for 3 weeks and normalized to atmospheric oxygen levels (n=3 measurements in one BM Chip compared to n=1 in the well culture).

**Supplementary Fig. 3.**
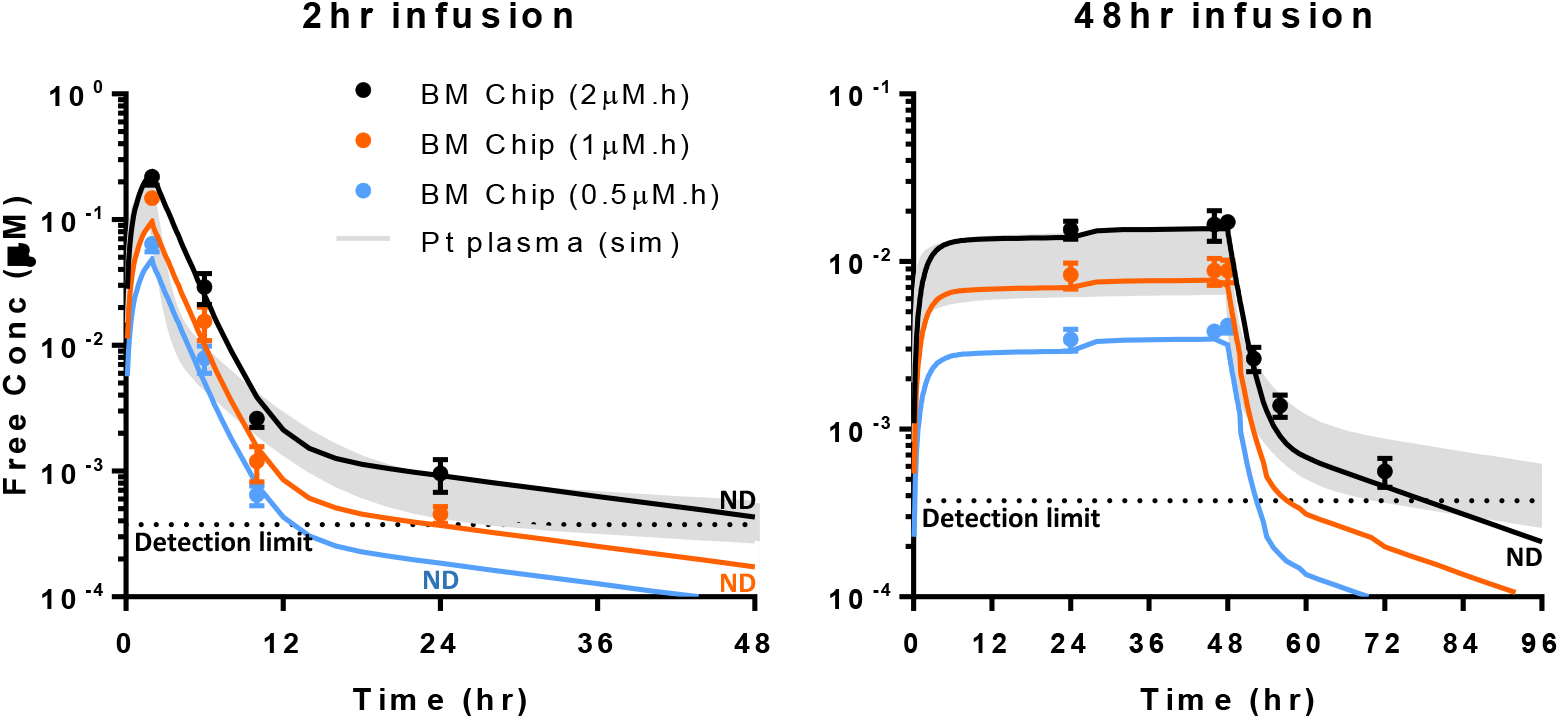
Results of PK models of BM Chip drug exposure (lines) derived from the measured drug concentrations for BM Chips (circles) that were matured for 10 days, treated with 2-hour or 48-hour infusions of AZD2811 (total AUC of 0.5μM.h, 1μM.h, and 2μM.h), and then cultured again in drug-free medium compared with an average patient’s plasma values that were simulated at a range of clinical doses^24^ based upon the known PK characteristics of AZD2811 (Pt plasma). Dotted line represents the detection limit of mass spectrometry during these experiments. AZD2811 concentrations were measured by mass spectrometry in BM Chip outlet medium collected at various timepoints during and after drug infusion (ND = not detectable).

**Supplementary Fig. 4.**
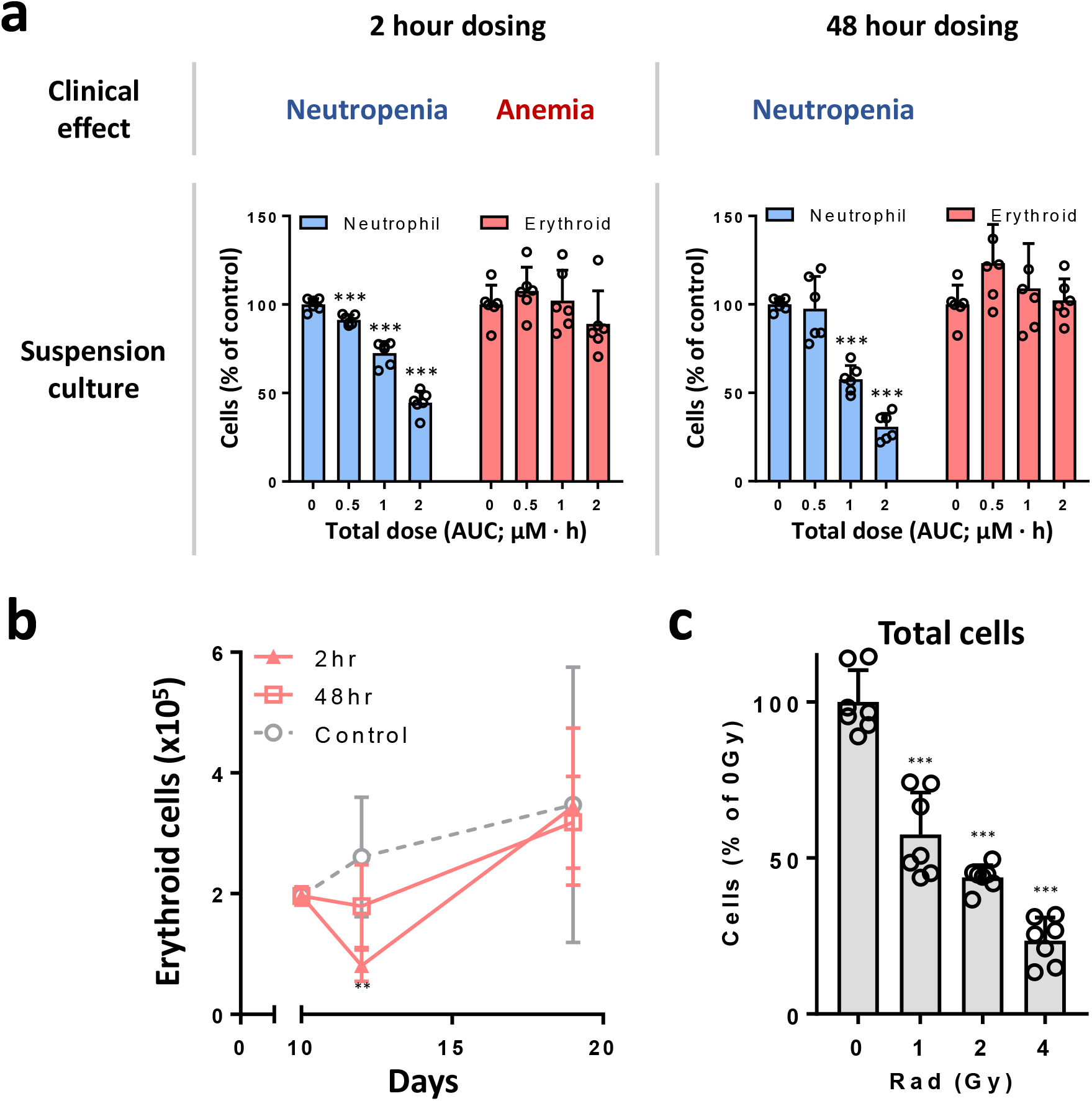
**a**, Hematologic side effects previously reported in patients receiving 2-hour or 48-hour infusions of similar total doses of AZD2811 administered as barasertib^23,24^ (top). Below are graphs showing effects of similar total doses of AZD2811 for 2-hours (left) versus 48-hours (right) on neutrophils (blue) and erythroid (red) cells when added to suspension cultures of CD34+ cells on day 10, harvesting the cells on day 12, and quantifying cell numbers using flow cytometry (n=6 wells per condition; data pooled from 2 independent experiments). **b**, Erythroid cell numbers quantified by flow cytometry from BM Chips that were matured for 10 days, treated with 2‐ or 48-hour infusions of the same total dose of AZD2811 (2μM.h), and then allowed to recover in drug-free medium. Analysis was performed at days 10, 12, and 19 (n=6 chips per condition at each timepoint; data pooled from 2 independent experiments). **c**, Total cell numbers in BM Chips that were matured for 10 days, irradiated with 0, 1, 2 or 4 Gy, and sacrificed at day 14 (n=7 chips, data pooled from 2 independent experiments; *P<0.05; **P<0.01; ***P<0.001).

**Supplementary Fig. 5.**
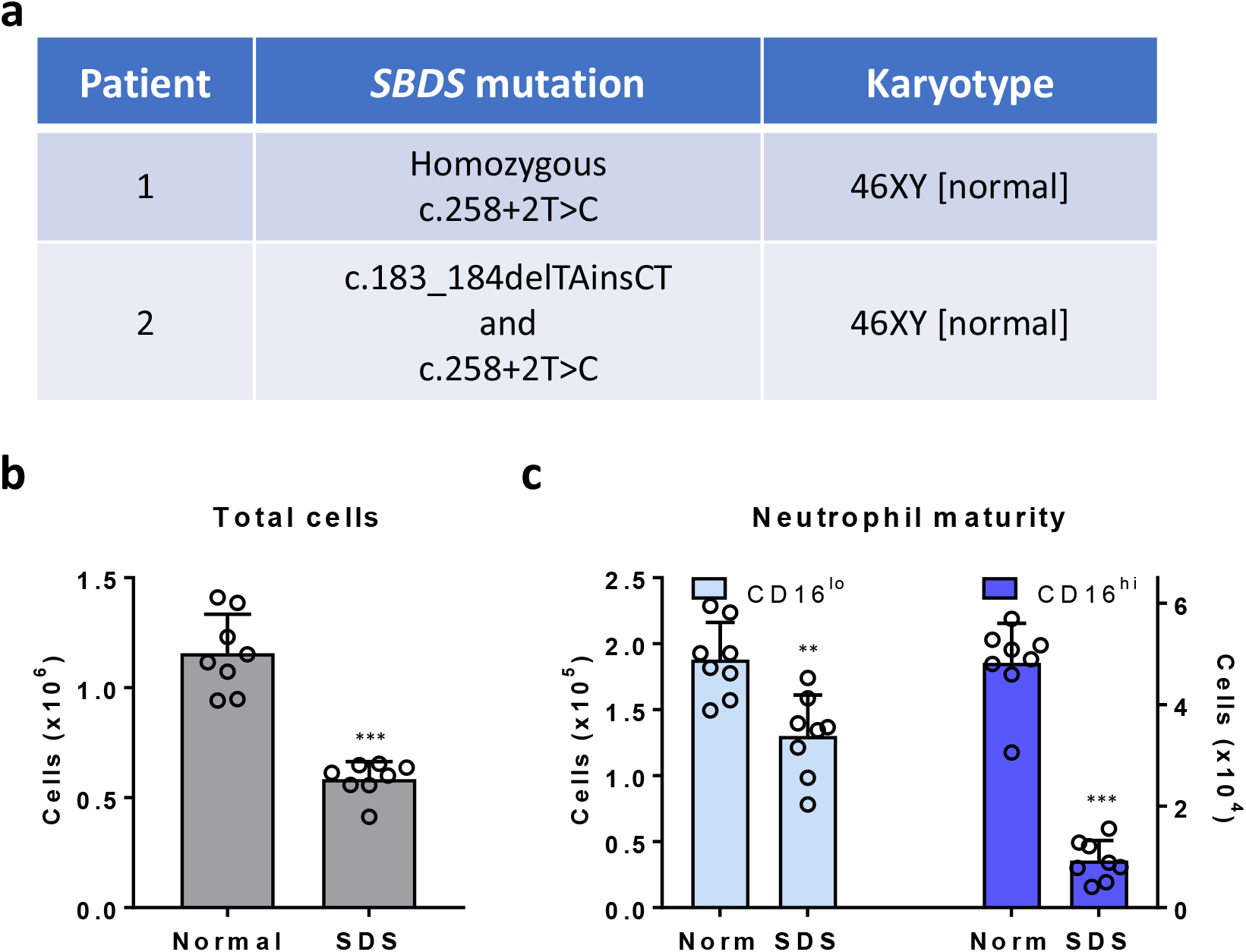
**a**, *SBDS* mutation characteristics and karyotype of the two SDS patients that provided cells for the BM Chips described in this study. **b,c**, BM Chips containing normal or SDS CD34+ cells that were matured for 14 days. Numbers of total cells (**b**) and immature CD16^lo^ vs mature CD16^hi^ neutrophils (**c**) were quantified by flow cytometry (n=8; pooled from 2 independent experiments using cells from different SDS patients with 4 chips per experiment; *P<0.05; **P<0.01; ***P<0.001).

**Supplementary Fig. 6.**
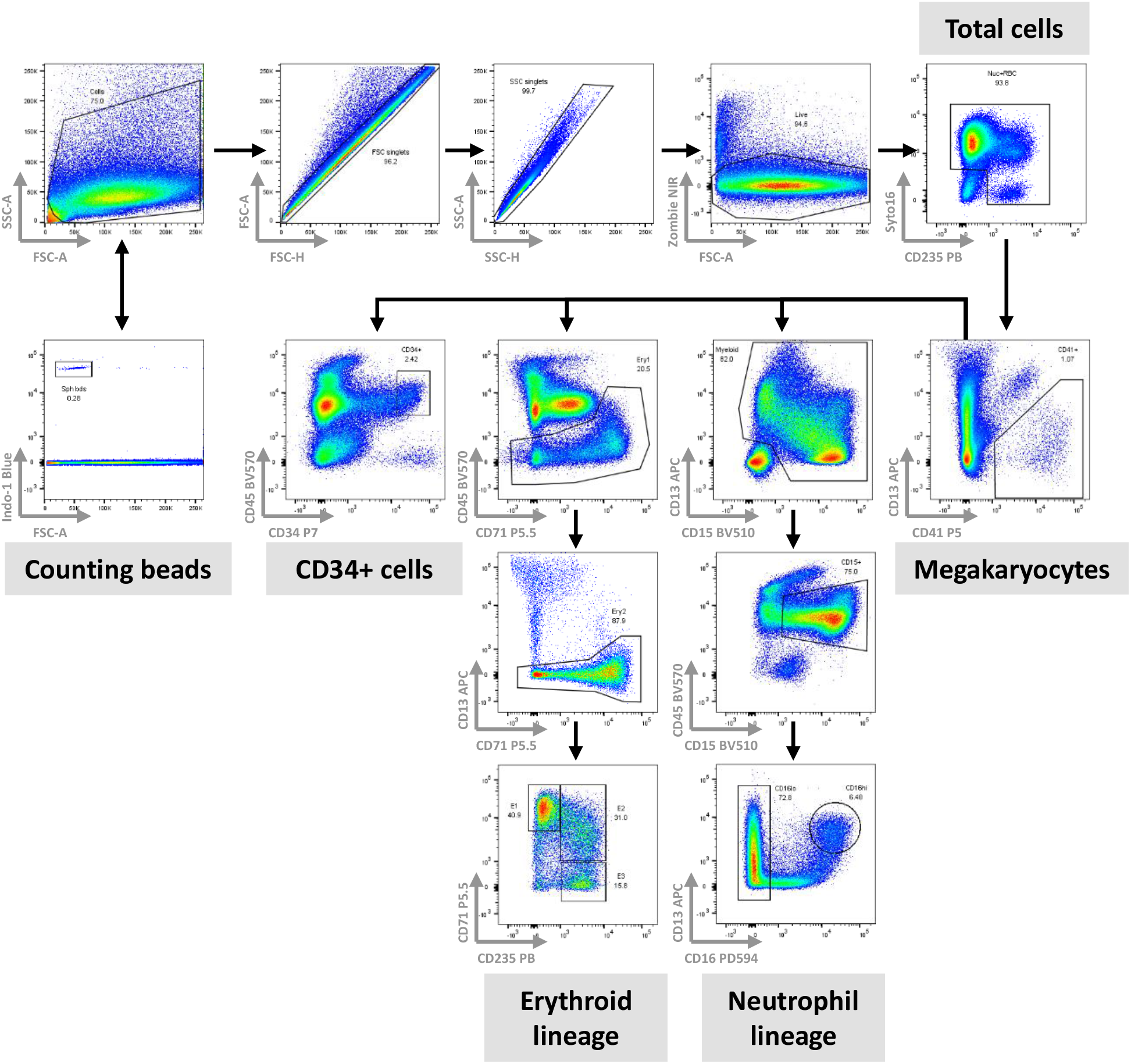
Representative flow cytometry gating scheme to quantify total, CD34+, neutrophil lineage, and erythroid lineage cells.

